# Alpha 9 integrin expression enables reconstruction of the spinal cord sensory pathway

**DOI:** 10.1101/2023.03.24.534172

**Authors:** Katerina Stepankova, Barbora Smejkalova, Lucia Machova Urdzikova, Katerina Haveliková, Fred de Winter, Stepanka Suchankova, Joost Verhaagen, Vit Herynek, Rostislav Turecek, Jessica Kwok, James Fawcett, Pavla Jendelova

**Author notes:** Corresponding address: Institute of Experimental Medicine, CAS Videnska 1083 142 00 Prague 4 Czechia.

## Abstract

Full recovery from spinal cord injury can only occur if the axon pathways connecting the brain and spinal cord regenerate and restore motor and sensory connections. Neither sensory nor motor axons can regenerate spontaneously in the spinal cord in mammals. This failure is partly due to the lack of suitable adhesion molecules on the sensory axons that allows them to interact with the environment of the damaged spinal cord. In this rat study, an integrin adhesion molecule along with its activator was expressed in sensory neurons using an adeno-associated viral (AAV) vector. Expression of these adhesion molecules allowed sensory axons to regenerate through the spinal cord injury and all the way back to the brainstem, restoring the sensory pathway. Treated animals regained touch sensation and sensory behaviours.

The integrin ligands in the injured spinal cord are tenascin-C and osteopontin, but adult PNS and CNS neurons lack receptors to them. Sensory neurons were transduced with α9 integrin, which combines with endogenous β1 as α9β1 (which is a tenascin/osteopontin receptor) together with the integrin activator kindlin-1. Regeneration from sensory neurons transduced with α9integrin and kindlin-1 was examined after C4 and after T10 dorsal column lesions with C6,7 and L4.5 sensory ganglia injected with AAV1 vectors. In animals treated with α9 integrin and kindlin-1, sensory axons regenerated through tenascin-C-expressing connective tissue strands and bridges across the lesion and then re-entered the CNS tissue.

Many axons regenerated rostrally to the level of the medulla. Regenerated axons were particularly visible at the border between white and grey matter in the dorsal cord. Stimulation of the median/sciatic nerve caused many neurons rostral to the injury to activate and express cFos. VGLUT1/2 staining indicated newly formed functional synapses above the lesion. Behavioural recovery was seen in heat, mechanical sensation and tape removal tests. Many axons regenerated from the thoracic lesions to the brainstem, a distance of 4-5 cm, equivalent to the length of 1 or 2 spinal segments in humans.

## Introduction

Sensory neurons in dorsal root ganglia (DRGs) send connections via peripheral nerves (PNS) and through the dorsal root into the spinal cord. When peripheral nerves are crushed many of the sensory axons regenerate back to skin and muscles, while damaged central branch axons show only a small regenerative response. After peripheral axotomy, DRG neurons express a genetic programme including many regeneration-associated genes (RAGs) ^1–3^ and the Schwann cells transform to a regeneration-permissive state ^4^. In this permissive state the Schwann cell surface contains axon growth-promoting extracellular matrix (ECM) molecules and various growth-promoting trophic factors are secreted. After sensory axons are damaged following spinal cord injury these events do not occur: the RAGs programme is not upregulated, and the glia in the cord and around the injury do not provide an environment that is permissive to axon growth. In particular, the parenchyma of the damaged cord does not provide ECM ligands that are recognized by sensory neuron integrins ^5–7^. Cell migration events such as axon regeneration are dependent on ligands in the environment matched to receptors in the cell ^8^. In a previous study DRG neurons were transduced with an integrin matched to the CNS environment. α9 integrin is a migration-inducing receptor for tenascin-C, an ECM integrin ligand which is present throughout the CNS ECM and upregulated around injuries ^9,10^. Additionally, the integrin activator kindlin-1 was co-transduced into the neurons to prevent inactivation of the integrins by inhibitory chondroitin sulphate proteoglycans (CSPGs) and NogoA ^11^. When these transduced neurons were axotomized in the dorsal root, sensory axons were able to regenerate back into the spinal cord and through the cord up to the level of the medulla ^12^. A subsequent study to profile mRNA changes in these transduced and regenerating neurons revealed that transduction with α9 integrin and kindlin-1 (α9-K1) upregulated the neuronal RAGs programme, and that axotomy and regeneration additionally upregulated a distinctive CNS regeneration programme that included many genes associated with autophagy, ubiquitination, protein degradation and axonal transport ^13^. Transduction with α9-K1 therefore provides to sensory neurons axotomized in the CNS the mechanisms that enable regeneration after PNS lesions, namely expression of the RAGs programme and an adhesion molecule matched to the environment. In the current study we have transduced sensory neurons with α9-K1 at the same time as performing dorsal spinal cord injuries at either C4 or T10 levels. This study investigates the ability of the transduced axotomized axons to traverse the non-permissive complex cellular environment of the damaged spinal cord, which includes reactive glia, fibroblast-like cells derived from the meninges, blood vessels, pericytes and inflammatory cells. The ability of axons that cross the lesion to regenerate up the spinal cord is then studied. We show that α9-K1-transduced axons regenerate vigorously through the spinal cord injury core, associating with cells that express tenascin-C and are then able to re-enter the CNS environment where they can regenerate as far as the medulla. The regenerated axons send side-branches into the dorsal horn of the spinal cord where they stimulate propriospinal neurons, and mediate full recovery of sensory functions.

## Materials and methods

### Preparation of AAV1 vectors

The plasmids AAV-SYN-α9-V5, and AAV-CMV-kindlin1-GFP were scaled and sequenced before proceeding to be packaged into AAV1 as described previously ^14^. For AAV vector production, HEK293T cells were transfected with the individual expression and helper plasmids and were cultured for 3 days. The transfected cells were lysed by using 3 freeze-thaw cycles. After centrifugation, the crude lysate was subjected to iodixanol gradient (15%, 25%, 40% and 60%) ultracentrifugation using a Type 70Ti rotor (Beckman) at 490000 × *g*, 16°C for 70 min. The AAV vector particles was collected and concentrated using an Amicon Ultra-15 device (Millipore). The titer of the virus was determined by using real-time quantitative PCR resulting in the following titers: 2.34 × 10^12^ GC/ml for AAV1-α9-V5 and 4.99 × 10^12^ GC/ml for AAV1-kindlin1-GFP. AAV1-SYN-GFP was purchased from Vigene (distributed by Charles River). In order to maintain the same injected titer for the experimental and control groups, the purchased AAV1-SYN-GFP was diluted from 2.0 × 1013 GC/ml to 2.0 × 1012 GC/ml.

### Experimental animals and animal surgeries

Seventy-two female Lister-Hooded rats (150-175 g; Envigo) were involved in the present study. Rats were housed three per cage on a 12h/12h light/dark cycle under standard conditions: temperature (22 ± 2°C) and humidity (50% ± 5%). The rats had free access to water and food ad libitum.

All procedures were approved by the ethical committee of the Institute of Experimental medicine of ASCR and performed in accordance with Law No. 77/2004 of the Czech Republic. Based on previous studies, the number of animals had been statistically optimized for each particular experiment to achieve their reduction according to European Commission Directive 2010/63/EU, and all efforts were made to minimize pain and suffering. Each animal was allocated a number and randomly assigned to one of the control or experimental groups. Experimenters were blinded throughout the entire study, including behavioural testing and quantification of axon growth. The identity of the animals and their treatment group was not revealed until after evaluation. The behavioural tests were conducted by two independent experimenters.

In order to compare the forelimb and hindlimb sensory regeneration after viral vector administration, the animals were randomly divided into 2 cohorts (n=36 per cohort). One cohort underwent the C4 level of injury with C6 and C7 DRG injections (for forelimb sensory functions restoration). The second cohort underwent the T10 lesion with L4 and L5 DRG injections (for hindlimb sensory functions restoration) (Fig. 1B).

**Figure 1.**
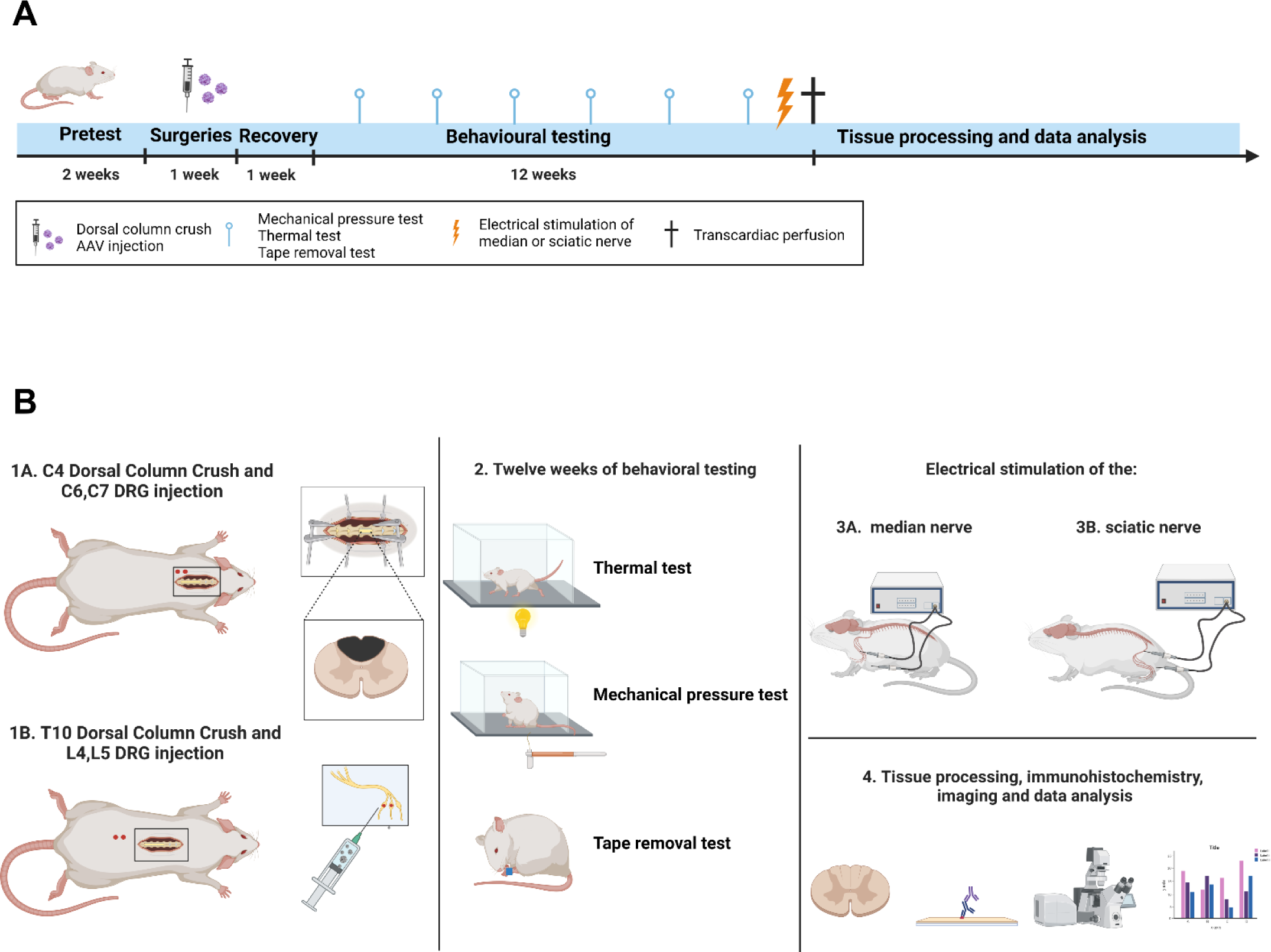
Experimental scheme. (**A**) Schematic illustration of the experimental timeline. (**B**) Diagram representing the experimental protocol. Both illustrations were created with BioRender.com.

Surgical procedures were performed under an inhalation anaesthesia with isoflurane (1.8-2.2%; Baxter Healthcare Pty Ltd; #26675-46-7) in 0.3 l/min oxygen and 0.6 l/min air. All animals received buprenorphine (Vetergesic® Multidose) subcutaneously at a dose of 0.2 mg/kg body weight. Using sterilized surgical equipment, two DRGs (C6, C7 or L4, L5 depending on the cohort) on the left side were exposed and 1 μl of viral vector per DRG at a working titre of 2 × 10^12^ GC/ml was manually injected using Hamilton syringe with custom-made needles (Hamilton; specification: 33 gauge, 12mm, PST3). At the same time, a concurrent laminectomy at the C4 or T10 level (depending on the cohort) was performed. A small slit in the dura was pierced with U100 insulin syringe (B Braun Omnican 50 Insulin Syringes; #9151117S), and the dorsal columns were crushed with a pair of fine Bonn forceps (Fine Science Tools). The tips of the Bonn forceps were held on either side of the dorsal columns at the site of the small slits, pushed 1 mm down into the spinal cord, and then held tightly for 15s. This lesion has been shown in many previous studies to cause a complete transection of the dorsal columns down to the level of the spinal canal. Immediately after the injury, muscles and skin were sutured. At the end of the experiment, the animals were intraperitoneally anesthetized with a lethal dose of ketamine (100 mg/kg) and xylazine (20 mg/kg) and perfused intracardially with 4% paraformaldehyde (PFA) in 1X-PBS. Two animals (one from each cohort) died the night after the surgery. We had to humanely euthanize two animals from the thoracic injury cohort for welfare reasons in this study.

### Sensory behavioural testing

The mechanical pressure (von Frey test) test, the thermal pain test (Hargreaves test), and the tape removal test were performed every other week for 12 weeks to ensure that animals did not learn the task independently of sensory reflexes. Animals were placed in the testing room at least 30 minutes before the test to allow adaption. Each rat was measured 3 times before the surgical procedure to obtain a baseline for each test.

### Mechanical pressure test

The touch sensitivity was measured with the electronic von Frey test (IITC Inc., Life Science Instruments, Woodland Hills, CA, USA). Animals were placed in the plexiglass enclosure on a mesh floor stand (IITC Inc., Life Science Instruments, Woodland Hills, CA, USA) at least 30 minutes before the test to allow adaption. Paws were then stimulated by a slowly-raising probe with rigid plastic tip to touch the centre of the paw. Pressure was increased until the nociceptive response of paw withdrawal. Thereafter, the value was recorded. Each paw was measured 5 times, forelimb (cervical SCI cohort) / hindlimb (thoracic SCI cohort), left (experimental) and right (internal control). The lowest and the highest of these five values were deleted and the three remaining values were averaged.

### Thermal test

SCI-mediated changes in thermal sensation were measured by the Ugo Basile Plantar Heat test apparatus (Comerio VA, Italy). Rats were placed into a plexiglass box with fiberglass bottom and let habituated there for 30 minutes. An infrared emitting device (Ugo Basile) was then placed directly under the footpad of the forelimb/hindlimb. The time when the rat withdrew its paw was recorded. The heat stimulus and timer were automatically activated simultaneously. The infrared stimulus turned off automatically after 30 s to prevent any harm to the animal. Five trials were performed on each forelimb (cervical SCI cohort) / hindlimb (thoracic SCI cohort), left (experimental) and right (internal control) with at least 3 min pause between individual trials. The average withdrawal time was calculated by averaging the three trials after deleting the lowest and the highest value for each animal.

### Tape removal test

For the tape removal test, each rat was trained over 5 sessions and then tested every other week post-surgery. The rat was then placed in an empty test cage and habituated there for 15 minutes. A small piece of tape (approximately 1 square centimetre) was taped to the paw. Three trials were performed on each forelimb (cervical SCI cohort) / hindlimb (thoracic SCI cohort), left (experimental) and right (internal control) with at least 3 min pause between each trial. Left and right paws were tested simultaneously. The time when the animal first noticed the tape and the time when animal removed the tape was recorded. If the rat did not notice/remove the tape by 5 minutes the tape was removed and a time of 5 min was recorded. For this test the paper tape (TimeMed Labeling Systems, Inc.; Fisher Scientific; USA; #NC9972972) was used.

### Electrostimulation

Three randomly selected animals from each group were terminally anaesthetised with 1.5g/kg urethane (Sigma Aldrich; #U2500). Then median nerve (cervical SCI cohort) or sciatic nerve (thoracic SCI cohort) was exposed. The stimulating electrode was inserted into the nerve and the ground electrode into the nearby muscle. The stimulation consisted of 10 stimulus trains, with stimulus duration of 0.5 ms with amplitude of 7.2 mA, a frequency of 100 Hz, and train duration of 2 s and 8 s intervals between trains as previously described by Bojovic et al ^15^. Used electrodes were single subdermal needles (Rhythmlink). Animals were perfused 2 hours after electrical stimulation.

### MRI

After transcardial perfusion, 2 cm long injured spinal cord samples were collected, post-fixed in 4% PFA in PBS (over 2 nights), then transferred to PBS with 0.002% azide in a small plastic tube (1.5 ml) with a cap. Spinal cords were then visualised ex vivo using a 7T preclinical MRI scanner (MRS*DRYMAG 7.0T, MR Solutions, Guildford, UK) equipped with a mouse head resonator coil. Three high resolution sequences were acquired: A T2-weighted turbo spin echo sequence in the axial direction, repetition time *TR* = 4000 ms, turbo factor *TF* = 8, echo spacing *TE* = 8 ms, effective *TE* = 40 ms, number of acquisitions *AC* = 16, acquisition time approximately 17 minutes. Acquired matrix was 128 × 128, field of view FOV = 10 x 10 mm^2^, 20 slices, slice thickness 0.5 mm with no gap. A similar T2-weighted turbo spin echo sequence was used for sagittal slices (*TR* = 3000 ms, *TF* = 8, *TE* = 8 ms, effective *TE* = 40 ms, *AC* = 16, acquisition time 13 minutes) with modified geometry: matrix 128 × 256, FOV = 10 × 20 mm^2^, 15 slices, slice thickness = 0.3 mm without gap. A T1-weighted axial image was acquired using a 3D gradient echo sequence, *TR* = 10 ms, flip angle 20°, *TE* = 3.3 ms, number of acquisitions *AC* = 16, acquisition time 11 minutes. The matrix was 128 × 128 × 32, field of view FOV = 10 × 10 × 16 mm^3^ (providing axial slices with a slice thickness of 0.5 mm). The MRI was performed in collaboration with the Centre for Advanced Preclinical Imaging (CAPI) in Prague.

### Immunostaining

PFA-fixed tissue samples were cryopreserved in sucrose solution with gradually increasing concentration (10%, 20%, and 30% sucrose in deionized water). Samples were transferred from a less concentrated solution to a more concentrated solution after immersion. Tissue was then embedded in OCT mounting media (VWR, #03820168). Sections were cut on slides at 8 μm (DRGs) and at 20 μm (spinal cords) on a cryostat (Thermo Scientific, Cryostar NX70). Sections were permeabilized in 0.5% Triton X-100 (Sigma, #T8787) for 2 h at room temperature (RT) and then blocked in 0.2% Triton X-100 and 10% ChemiBLOCKER (Millipore, #2170) for another 2 h at RT. After blocking, sections were incubated with following primary antibodies and/or labelling agents: anti-GFP (Invitrogen, #11122, 1:800, 3 days or Invitrogen, #10262, 1:800, 3 days), anti-V5 (Invitrogen, #R96025, 1:800, 3 days), anti-cFOS (Abcam, #ab208942, 1:500, 2 days), anti-neurofilament (NF200) (Sigma, #N4142, 1:800, 3 days), anti-IBA4 (Sigma, #L2140, 1:800, 3 days), anti-CGRP (Sigma, #PC205L, 1:800, 3 days), anti-β-III tubulin (Cell Signaling, #5568S, 1:1000, 3 days), anti-laminin (Abcam, #ab11575, 1:800, 3 days), anti-tenascin-C (obtained from Faissner lab, Bochum, Germany, 1:500, 2 days), anti-GFAP-Cy3 (Sigma-Aldrich, #C9205, 1:800, 3 days), and anti-VGLUT1/2 (Synaptic Systems, #1235503, 1:800, 3 days). Goat anti-host antibodies of the respective primary antibodies conjugated with Alexa Fluor 405, 488, 594 and, 647, (1:300; 4 h, RT, Invitrogen) were used as secondary antibodies. Tissues were then washed with 0.2% Triton X-100 in PBS and subsequently mounted with Mowiol mounting medium (Carl Roth, #0713.2) with added DABCO to reduce fluorescent signal fading (Carl Roth, #0718.1).

### Tissue Clearing for light sheet microscopy

Four randomly chosen DRGs from each group were dehydrated by using ethanol (EtOH) dilution series (30%, 50%, 70%, 100%) with 2% Tween20 (Sigma-Aldrich, #P1379) and then delipidated in DCM with EtOH (2:1). Samples were then rehydrated by serial EtOH diluted solutions (70%, 50%, 30%) with 2% Tween20. Samples were then permeabilized by solution containing 0.2% triton v PBS, 0.3M glycine, 10% DMSO. Samples were blocked in 0.2% triton v PBS, 0.3M glycine, 10% DMSO, and 10% ChemiBLOCKER (2 days, 37°C). After blocking, DRG samples were incubated in the blocking solution with added primary antibodies: anti-GFP (Invitrogen, #11122, 1:400, 2 days, 37°C), and anti-V5 (Invitrogen, #R96025, 1:400, 2 days, 37°C). Secondary antibodies (1:300) were prepared in blocking solution and applied for 3 days at 37°C. Samples were then dehydrated again by using the same EtOH dilution series as at the beginning, and then cleared in Ethyl Cinnamate (Sigma-Aldrich, #112372) for 2 hours at RT. After clearing and imaging were performed, the DRGs were rehydrated in EtOH diluted solutions (100%, 70%, 50%, 30%), washed in PBS and cryopreserved in sucrose solution (10%, 20%, 30%), sectioned as described previously, and re-used for transduction efficiency analysis.

### Microscopy

Confocal imaging was performed using a Zeiss LSM880 Airyscan microscope and lightsheet imaging was performed using Zeiss Lightsheet Z.1 microscope. Images were analysed semiautomatically using NIH ImageJ or by manually using the microscope eyepiece reticle if necessary. Lightsheet data were processed by Huygens Software ((Scientific Volume Imaging, The Netherlands, http://svi.nl) and Imaris Microscopy Image Analysis Software (Oxford Instruments). The images used for the analysis were not manipulated, and the unedited data and metadata are archived by the authors. In the case of the submitted images, the curves were adjusted in all images and across the entire image. A philtre (unsharp mask) was used to make the axons more visible in the publication. Software used free and open-source raster graphics editor GIMP (The GIMP Development Team. (2019). GIMP. Retrieved from https://www.gimp.org).

### Statistical analysis

Data are expressed as mean ± SEM. Statistical differences between groups were determined by one-way or two-way with Tukey’s multiple comparisons test. For all statistical analyses, a *p* value of 0.05 was considered to be significant. Data processing and statistical analysis were performed using GraphPad Prism (GraphPad Software).

## Results

Female adult rats received spinal cord dorsal column lesions at level C4 or T9 and at the same operation received injections of AAVs to DRGs at level C6,7 or L3,4. The three experimental groups received different AAV injections, 1) AAV-GFP (GFP group), 2) AAV-kindlin-1-GFP (kindlin group), 3) AAV-kindlin-1-GFP + AAV-α9 –V5 (α9-K1 group). AAV-GFP (provides a control for AAV injection, AAV-kindlin-1-GFP shows the effect of activating the integrins that are endogenously expressed in DRG neurons, AAV-kindlin-1-GFP + AAV-α9 –V5 is the regeneration-inducing treatment with the tenascin-binding integrin and its activator. Animals received sensory behavioural testing for 12 weeks, then at week 13 animals were killed for histology. Before this some animals received electrical stimulation to their sciatic or median nerve to upregulate cFOS in spinal neurons (Fig. 1).

### AAVs transduce integrin and kindlin into DRG neurons

Transduction of DRG neurons was assessed by immunostaining of the DRGs and axons below the level of the lesions (Supplementary Fig. S1A). Transduction efficiencies were obtained by counting the ratio of GFP and/or V5-positive cell bodies with βIII-tubulin-positive cell bodies. Transduction efficiency was similar for the three vectors and ranged from 28-38% for single vectors and 20-25% for co-transduction with both α 9 and kindlin1 vectors. The kindlin1 vector was injected at a 1:3 ratio. Prior to the actual experiment, several ratios were tested (1:1; 1:3; 1:6). The 1:3 ratio gave the highest co-expression (data not shown). The transduction efficiency is shown in Table 1 and Fig. S1B To confirm that transduced sensory neurons are located throughout the DRG and not just locally, we imaged transduced DRGs using 3D light-sheet microscopy 13 weeks after AAVs injections (Supplementary Fig. S1C). In addition, immunostaining showed that AAV1-α9-V5 and AAV1–kindlin-1–GFP transduced all three sensory neuronal types located in DRGs: large diameter mechanoreceptors (NF200-positive), medium diameter nociceptors (IBA4-positive), and small diameter thermoreceptors (CGRP-positive) (Supplementary Fig. S1D).

**Table 1.** Transduction efficiencies of sensory neurons after direct DRG injection. Data are expressed as mean ± SEM, n= 20-24 per group.

| **VIRAL VECTORS** | **C6, C7 DRGs (%)** | **L4, L5 DRGs (%)** |
| --- | --- | --- |
| AAV1-GFP | 33.39 ± 1.714 | 36.59 ± 2.838 |
| AAV1-kindlin-1-GFP | 28.98 ± 0.780 | 39.06 ± 2.973 |
| AAV1-α9-V5 | 37.24 ± 2.194 | 32.94 ± 1.575 |
| % βIII tubulin +ve neurons co-transduced with AAV1-α9-V5 + AAV1-kindlin-1-GFP (injected in 3:1 ratio) | 21.75 ± 1.392 | 23.37 ± 1.589 |
| % α9 neurons also +ve for kindlin | 62.24 ± 4.570 | 68.48 ± 2.537 |

**Supplementary Figure S1.**
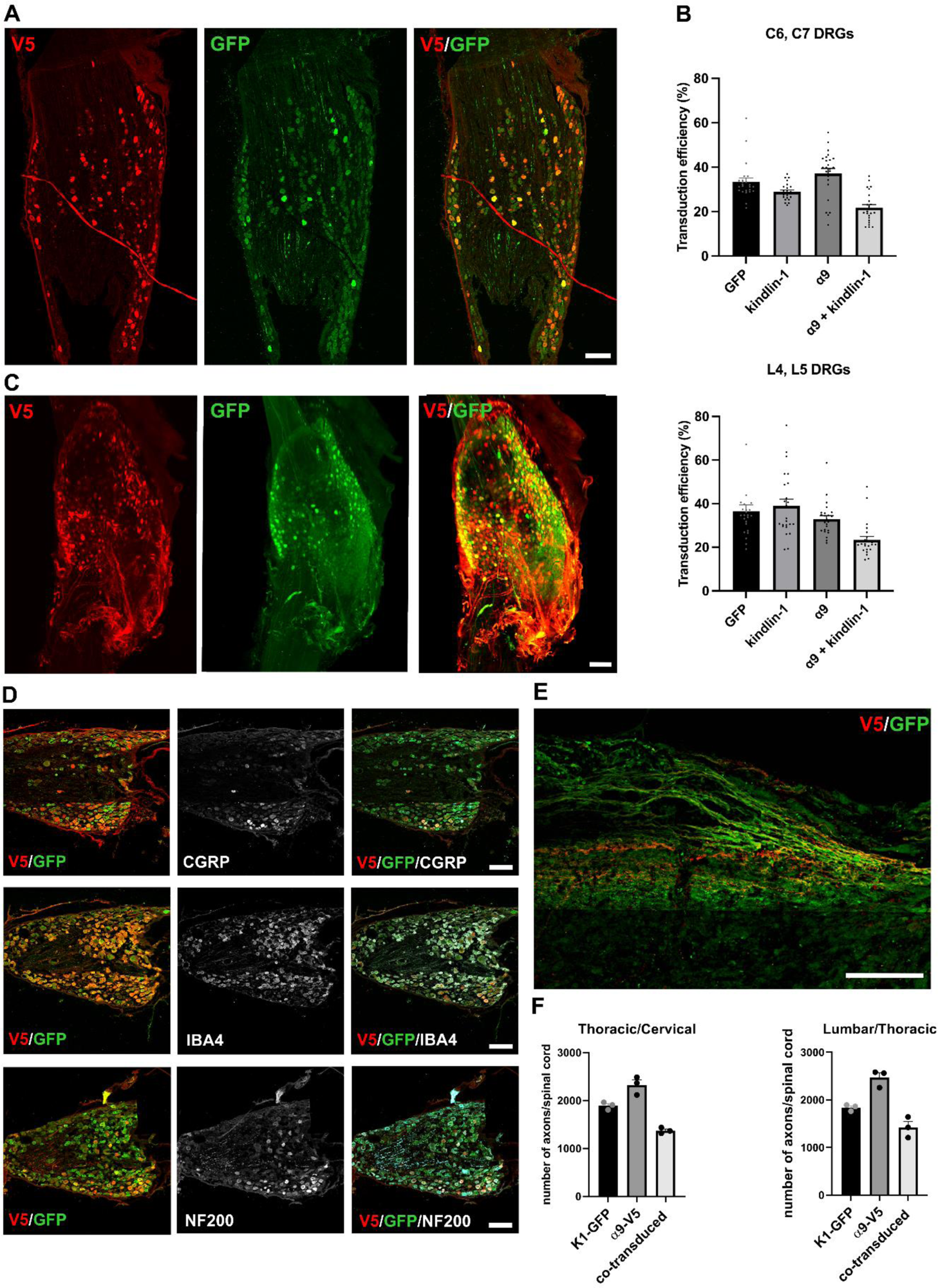
Expression of α9 integrin and kindlin-1 in DRG neurons and in the axons below lesion 13 weeks after direct injection. (**A**) A dorsal root ganglion that was injected 13 weeks ago with AAV1-kindlin-1-GFP + AAV1-α9–V5. Many neurons are yellow in the composite picture below, indicating co-transduction, but some neurons are red or green indicating that they were transduced by only one of the viral vectors. Scale bar: 200 μm. (**B**) Quantification of (A). (**C**) Light sheet microscope images showing the DRG after direct AAV injection with AAV1-kindlin-1-GFP + AAV1-α9-V5. Sensory neurons are uniformly transduced throughout the DRG. Scale bar: 200 μm. (**D**) Sensory neurons of the three main types were transduced with the AAV1 vectors. CGRP, IBA4 and NF200 immunostaining (middle column) overlaps with V5 and GFP staining showing integrin and kindlin (left and right composite columns). Scale bar: 200 μm (**E**) Axons caudal to the lesion stained for V5 and GFP. Many of the axons contain both integrin and kindlin, but there are axons that are only positive for one or the other. Scale bar: 100 μm. (**F**) Quantification of (**E**). Bar graphs show the individual data together with their mean ± SEM (n= 3 animals per groups).

The transport of α9-V5 and kindlin-1-GFP in sensory axons caudal to the lesion was visualized by immunostaining. The spinal cord was sectioned at 40µm then every fifth section was stained for V5 and GFP. These molecules transported from the DRGs were used as axon tracers for all the experiments in this study.

The number of axons was counted using an eyepiece grid, counting the number of axons crossing a line of the grid at regular distances caudal and rostral to the lesion. The sum of the values from each distance point was then multiplied by 5 to estimate the number of labelled axons at this level in each spinal cord.

In the dorsal roots as well as in the spinal cord caudal to the lesion, many axons were positive for both α9-V5 and kindlin-1-GFP, but there were also axons that were only positive for one molecule (Supplementary Fig. S1E,F). The observed average number of labelled co-transduced axons in the spinal cord below cervical lesion was between 1350 and 1550 for both, cervical and thoracic lesions (See Table 2).

**Table 2.** Number of labelled axons per spinal cord 600µm caudal to the lesion. Data are expressed as mean ± SEM, n= 3 per group.

|  | Cervical SCI cohort | Thoracic SCI cohort |
| --- | --- | --- |
| Kindlin-1-GFP axons | 1837 ± 39 | 1890 ± 45 |
| α9-V5 axons | 2465 ± 105 | 2322 ± 109 |
| Co-transduced axons | 1422 ± 124 | 1375 ± 35 |

### α9-K1 axons regenerate across lesions in connective tissue bridges

Axons from the integrin 9 + kindlin-1 group were observed to cross the lesion, re-enter CNS tissue and continue to grow rostrally up the cord. Within the lesions many axons were seen in GFAP-negative connective tissue strands and bridges and in the meningeal/connective tissue roof that covered most lesions, with a few regenerating axons growing around the lesion base particularly in cervical lesions. Where the connective tissue interfaced with the rostral lesion margin axons often showed tangles of axons with changes in course (Fig. 2 A,B), but once established in CNS tissue, axons followed a fairly straight course. Some axons did not re-enter CNS tissue but were seen growing alongside the cord in the meninges (Fig. 2C). In the α9-kindlin we measured 849 ± 64 axons 1mm rostral to the lesion margin.

**Figure 2.**
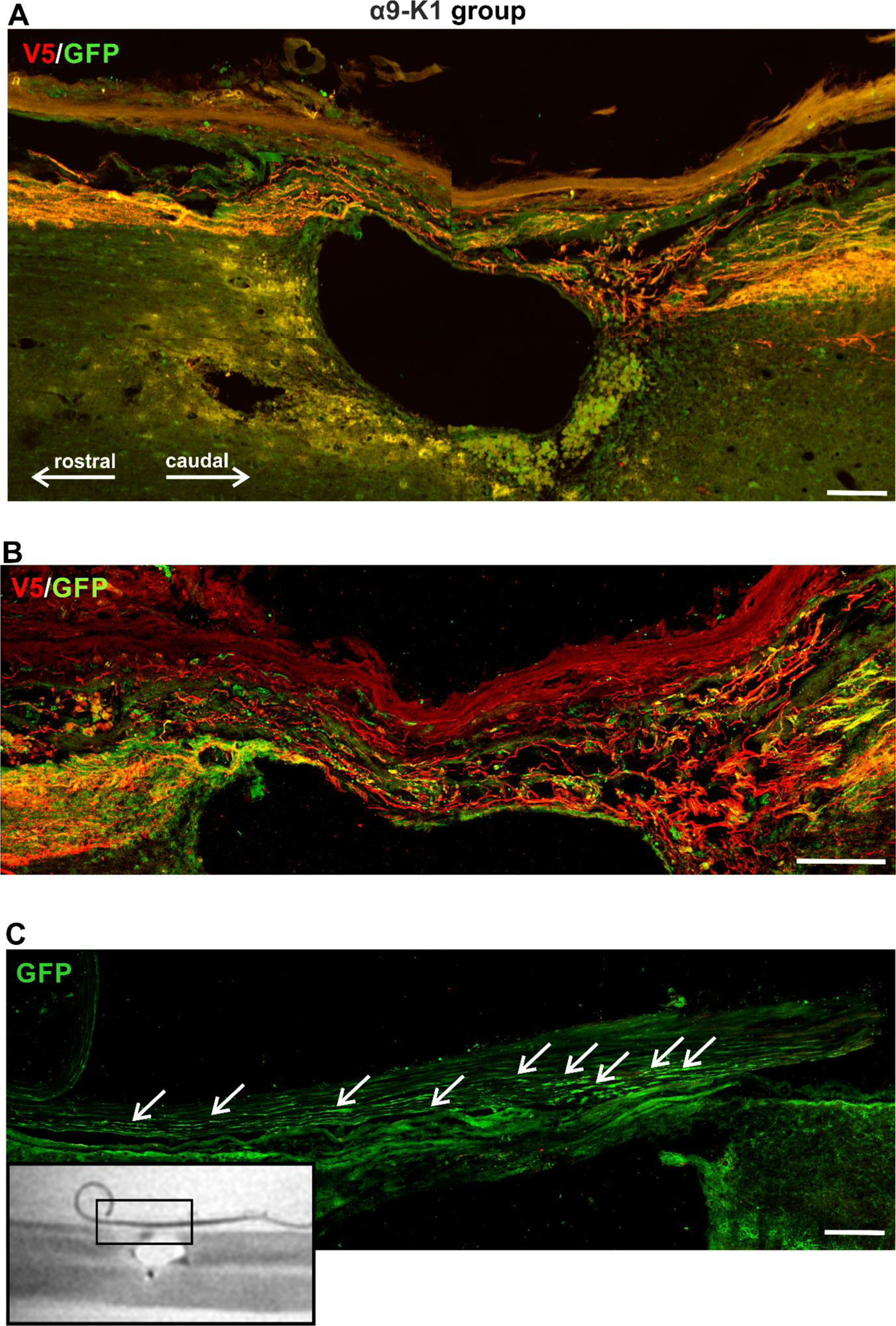
Co-expression of α9 integrin and kindlin-1 promotes sensory axon regeneration. (**A**) An example of spinal lesions at T10 from the α9-kindlin group. Many red (α9-V5 stained) axons approach the lesion from the right (caudal). There is some random growth as they enter the bridge across the top of the lesion that is mostly composed of meningeal cells, then the axons re-enter CNS tissue to grow rostrally on the left. Scale bar: 200 μm. (**B**) An example of axons passing through a fine connective tissue strand. At the rostral end (left) there is a region of wandering growth as the axons re-enter CNS tissue. Scale bar: 100 μm. (**C**) Where axons reach the point where connective tissue strands interface with CNS tissue, some axons continue to grow in the meninges besides CNS tissue (white arrows). The MRI picture on the bottom left shows where the detail came from. Scale bar: 50 μm.

In GFP controls no axons grew across the lesion, although a few sprouts were seen around the lesion edge and in the caudal margin of the lesion core (Fig. 3A-C). In animals expressing kindlin-1 alone some sensory axon regeneration was seen in the laminin+ve connective tissue within and bridging the lesion, closely in contact with laminin+ve structures, but axons did not re-enter CNS tissue to grow beyond the lesion area (Fig. 3D-F). On the margin of the rostral part of the lesion, the integrin α9-kindlin-1 group showed significantly more axons (819.00 ± 94) to the kindlin-1-only (413± 87; p = 0.0012, 2way-ANOVA, n = 10 – 12). However, no axons from kindlin-1-only group continued to grow rostrally above lesion and 1mm above lesion there were no axons visible neither from kindlin-1-only nor from GFP controls.

**Figure 3.**
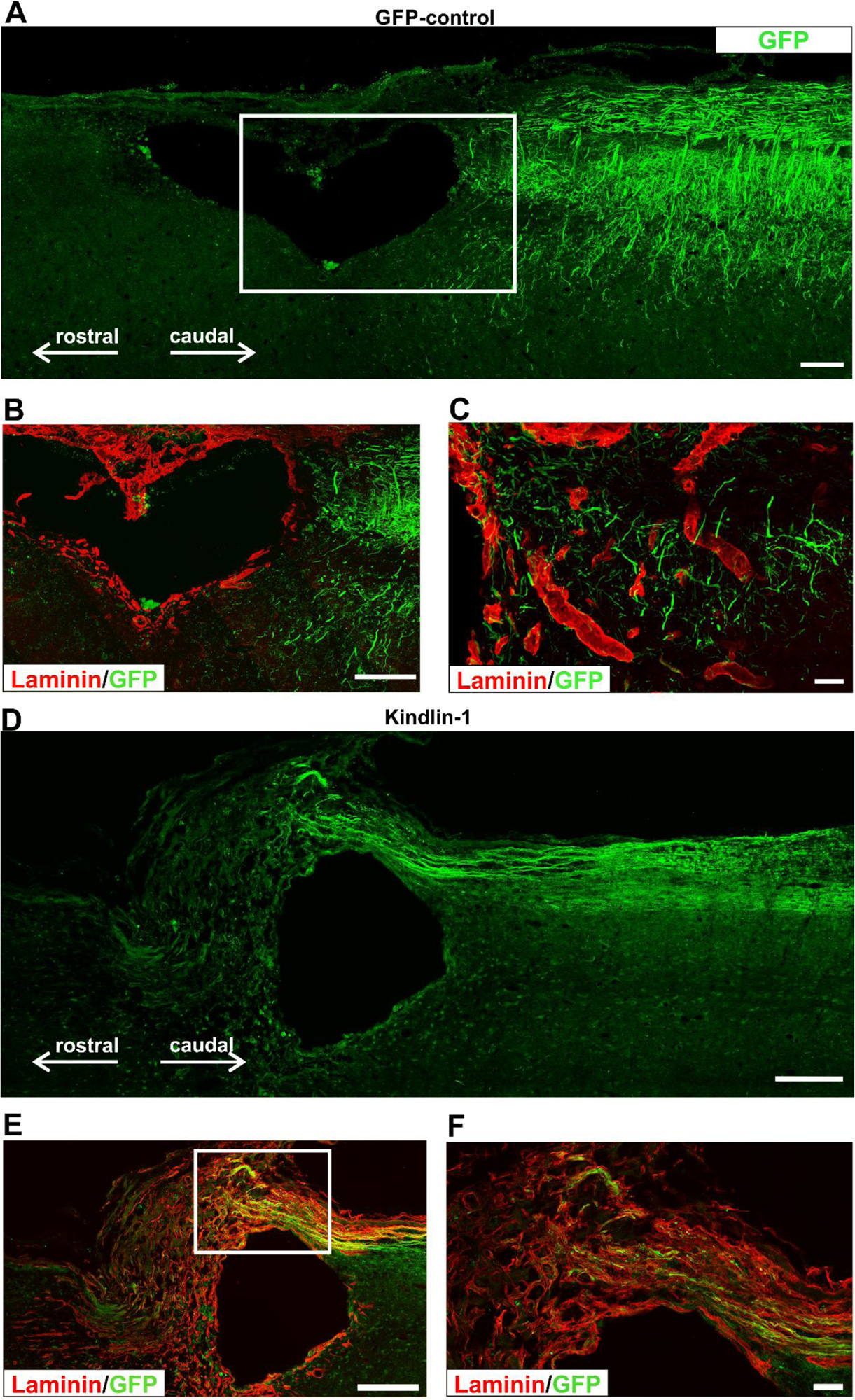
Axon growth was not observed in either the GFP or kindlin-1-only groups passing through the lesion. (**A**) A lesion from the GFP group immunostained for GFP. Axons can be seen only caudally to the lesion (right), but none have regenerated into bridge region that develops over lesion. Scale bar: 200 μm. (**B**) A lesion from the GFP group (inset from A), immunostained for GFP and laminin GFP+ve axons terminate caudally of the lesion (right), but none have regenerated into the laminin+ve connective tissue in the lesion. Scale bar: 200 μm. (**C**) A detail of the terminating axons of the GFP group where the laminin+ve boundary begins. The axons do not grow on the laminin+ve ECM. Scale bar: 50 μm. (**D**) Axon regeneration example from the kindlin-1 group. Scale bar: 200 μm. (**E**) Axons have regenerated into the laminin+ve connective tissue in the bridge region over the lesion, and are closely associated with laminin+ve processes. No axons have grown out of the connective tissue back into CNS tissue. Scale bar: 200 μm. (**F**) A magnified insets from (**E**). Scale bar: 50 μm.

### α9-K1 axons regenerate to the brain stem

Animals injected with both, integrin α9 integrin and kindlin-1 (α9-kindlin group), showed regenerating axons rostral to the lesion after both, C4 and T10 injury. Many of these were seen all the way up to the medulla. Most axons were seen at the interface between grey and white matter at the medial edge of the dorsal horns, and a lesser number of axons was seen following a wandering course within the dorsal horn white matter (Fig. 4A,B). These regenerating axons were therefore following a different pathway to that of unlesioned sensory axons. Along this regenerated pathway, there were frequent branches extending into the dorsal horn grey matter and terminating in arborizations (Fig. 4C). The distance over which axons regenerated from thoracic lesions reached up to 5cm. Almost all the regenerated axons rostral to the lesion stained for both α9-V5 and kindlin-1-GFP (Fig. 4D), in contrast to the axons caudal to the lesion, where there was a significant proportion of single-stained axons (Supplementary Fig. S1E). The implication is that only axons containing both α9 and kindlin-1 were able to regenerate through the lesion. Axon regenerative behaviour was similar in both cervical and thoracic lesions. The regeneration index 5 mm above lesion was approx. 0.5 for both, cervical and thoracic SCI and 0.2 in T10 SCI group 5 cm above lesion. Detailed information on the number of axons regenerating through C4 and T10 lesion is given in Table 3.

**Figure 4.**
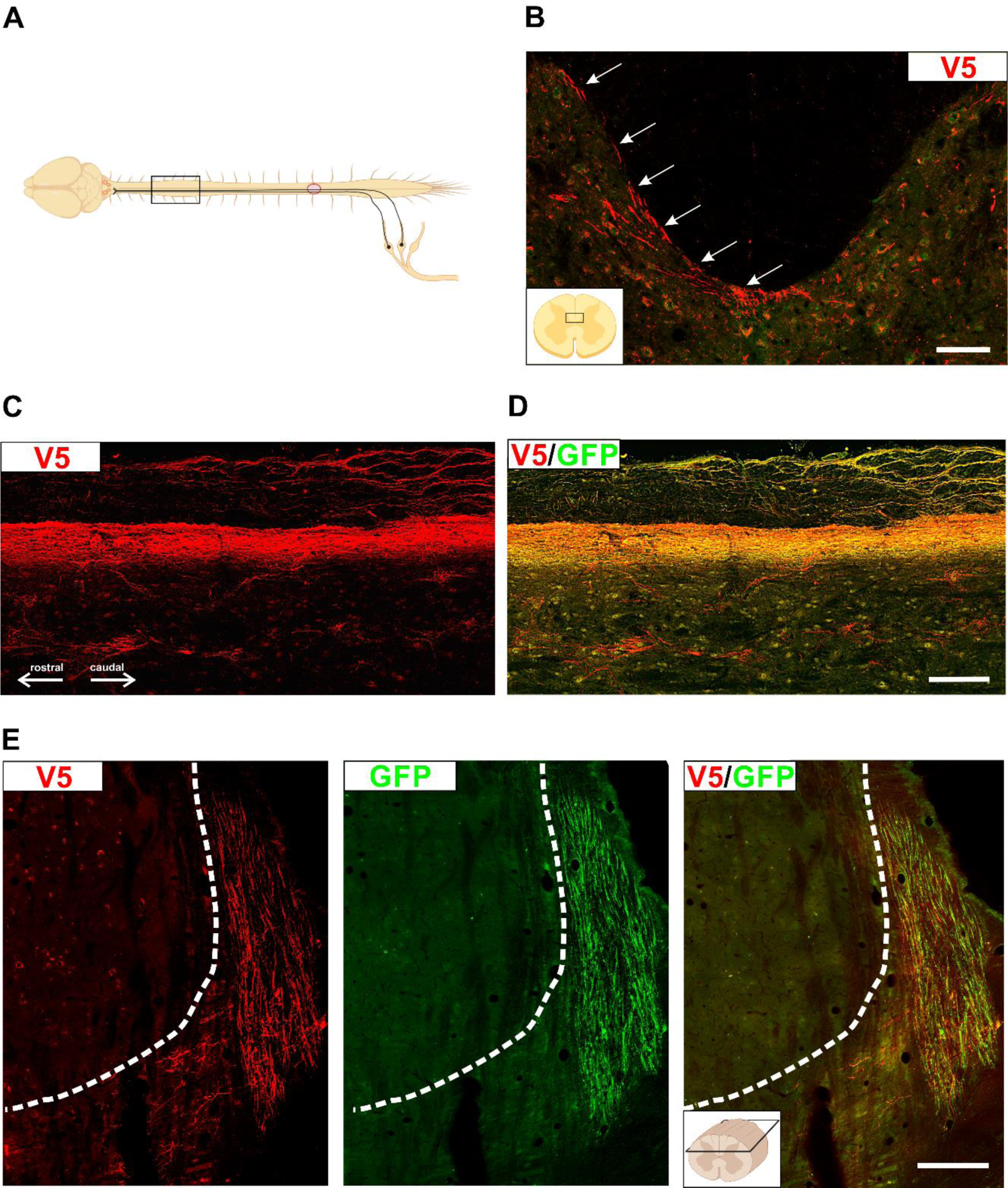
α9-K1 axons regenerate to the brain stem. (**A**) Diagram showing the spinal cord segment from which the following sections (**B, C, D**) were taken - an animal from the α9-kindlin group with a thoracic lesion and AAV injections into L4,5 DRGs. Created with BioRender.com. (**B**) A transverse section of the cervical cord (C3) with α9-V5 labelled axons. The diagram on the bottom left shows where the detail came from. The diagram was created with BioRender.com. Scale bar: 50 μm. White arrows point to the axons at the edge between the dorsal horn and the dorsal column. (**C**) Axons at level C3 labelled only for V5. The axons are following a pathway at the margin between the dorsal horn and the dorsal column with branches to the dorsal horn. Many axons are seen in a layer between dorsal horn and dorsal column white matter, and some axons are seen in the dorsal horn white matter. (**D**) Axons rostral to the lesion were double-stained for α9-V5 and kindlin-1-GFP, showing colocalisation of both markers. Scale bar: 100 μm. (**E**) Longitudinal sections medulla double stained for α9-V5 and kindlin-1-GFP with cuneate nucleus on top. The bundle of axons is approaching the edge of the nucleus and some are growing towards it but not into it. There are no labelled axons in the nucleus, indicating that there are no unlesioned sensory axons. Scale bar of longitudinal section: 100 μm.

**Table 3:**
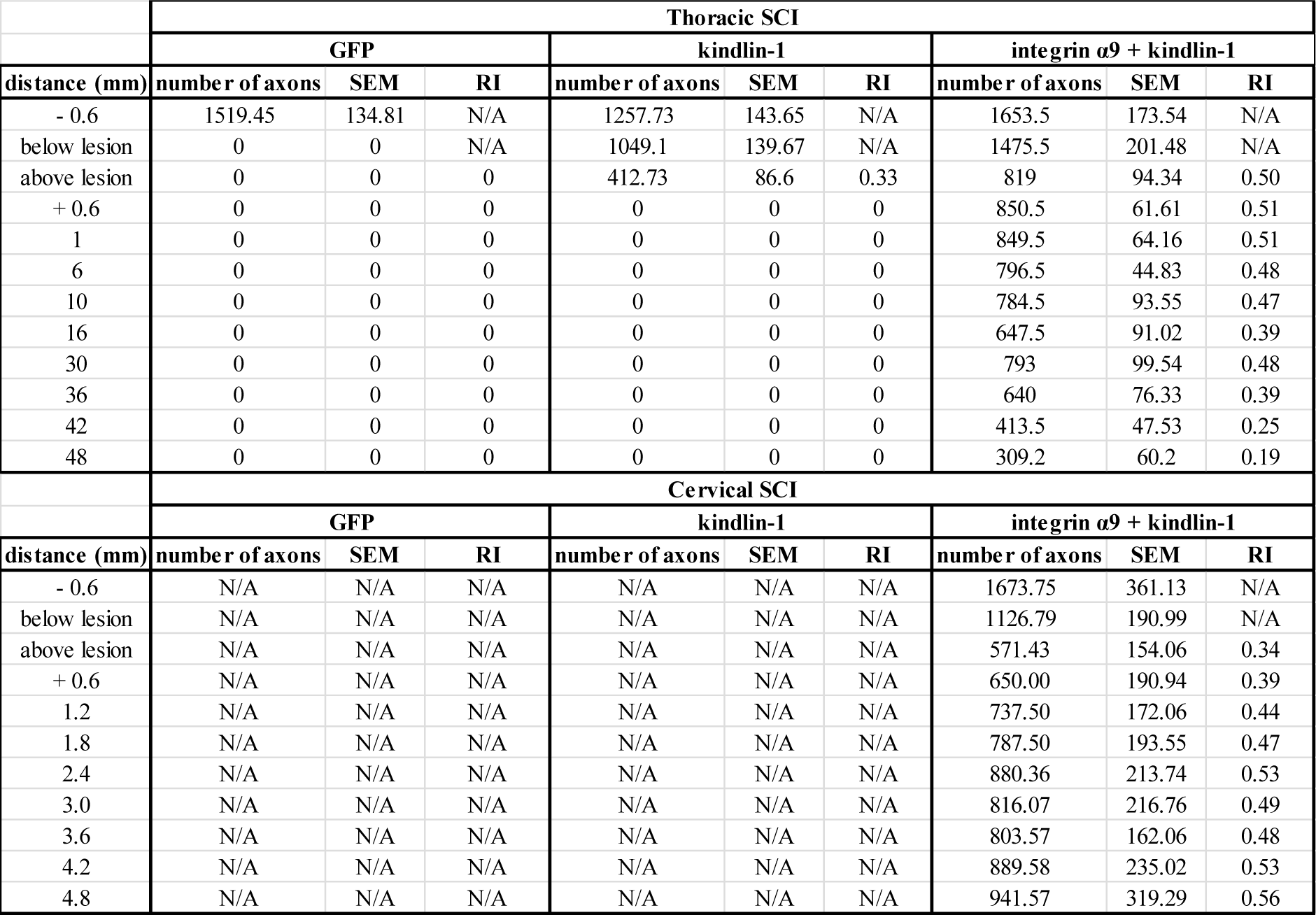
Number of axons and regeneration index in GFP, kindlin1 and α9-kindlin group. Data are expressed as mean ± SEM, n=7-10 (animals/group) NA=not available

As in previous studies ^12,16^ no α9-V5 or kindlin-1 +ve axons were seen innervating the two dorsal column sensory nuclei. Axons were seen around the margins of the nuclei but did not enter them (Fig. 4E). The absence of labelled axons in the nuclei confirms that the spinal cord lesions completely severed the sensory pathway. A summary of axon behaviour in the three experimental groups with axon counts for the α9-V5 or kindlin-1 +ve axons is shown in Fig. 5.

**Figure 5.**
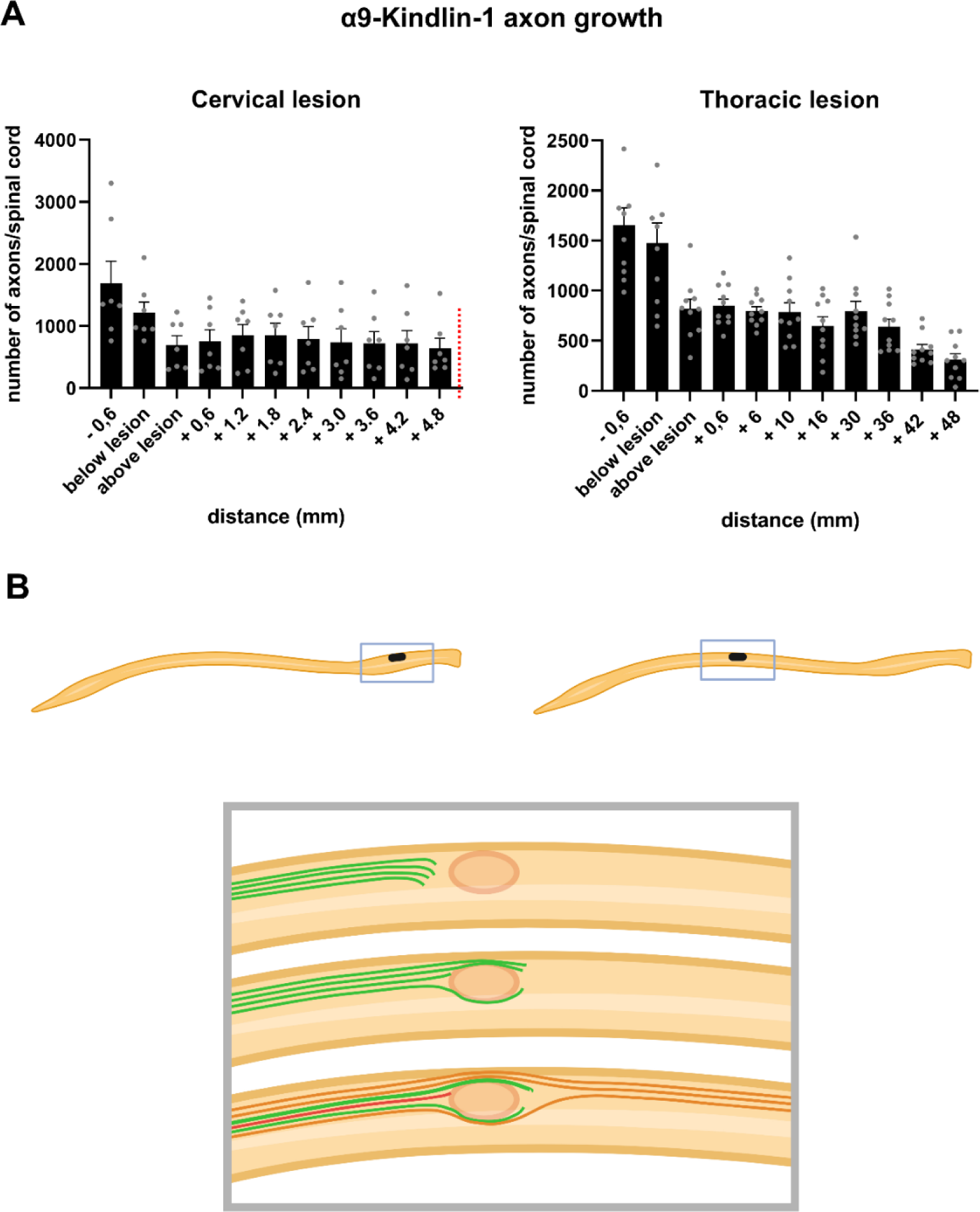
Distant regeneration in the spinal cord with co-expression of α9 integrin and kindlin-1. (**A**) Bar graphs show the number of axons after cervical injury with C6, C7 DRG injections (left) and thoracic injury with L4, L5 DRG injections (right). The red dotted line in the left graph indicates the counting limit, not the maximum distance the axons reached in the cervical SCI cohort. Approximately 900 sensory axons grew into the rostral spinal cord in the α9-kindlin group, with their number decreasing slightly as they approached the hindbrain. The distance of the thoracic lesion from the medulla is 5cm, where regenerated axons were observed. Bar graphs show data together with their means ± SEM (n= 7-12 animals per groups). (**B**) Schematic representation of the key results of this study. Only in the α9-kindlin group was there substantial regeneration beyond the lesion into the rostral cord. In the kindlin group axons regenerated in the laminin-containing connective tissue in the lesion core. Created with BioRender.com. No difference was observed between cervical and thoracic injuries in terms of the pathways followed by axons under different AAV-expressing conditions.

### α9-K1 axons regenerate through tenascin-C containing tissue

To examine the substrate on which α9-kindlin-1 V5-positive axons grew, immunolabeling for GFAP, laminin and tenascin-C was performed. The connective tissue strands and roof in lesions through which axons regenerated were entirely or partially GFAP negative, usually with a clear boundary with GFAP+ve CNS tissue. The connective tissue through which axons grew across the lesions stained for laminin and tenascin-C. The boundary between connective tissue and CNS was less clear with tenascin staining because the peri-lesional CNS tissue also expresses tenascin (Fig. 6A-C). In the kindlin-1 group GFP-positive regenerated axons group were observed in association with laminin +ve connective tissue substrate, but were not able to grow back into CNS tissue (Fig. 3D-F).

**Figure 6.**
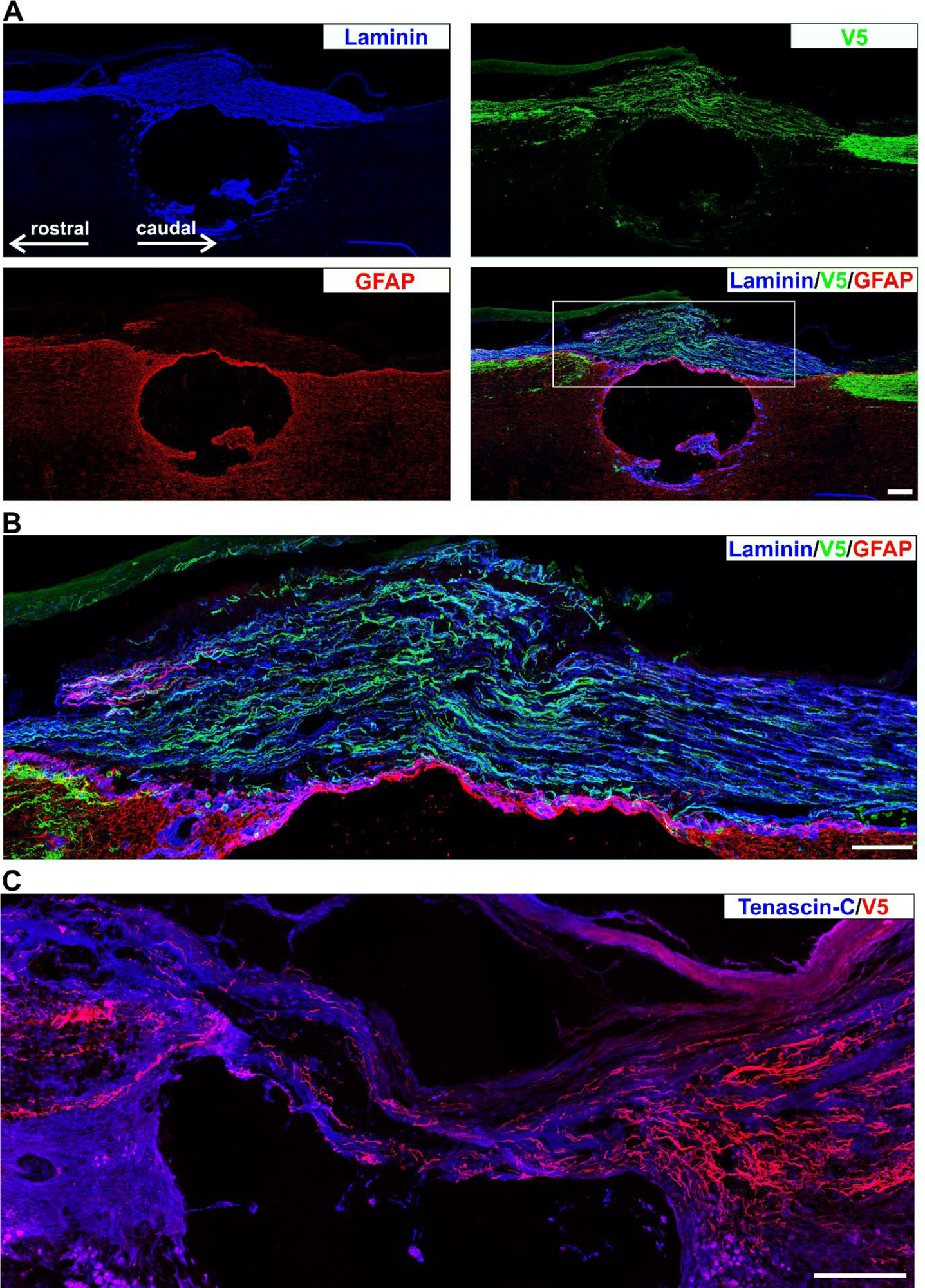
α9 integrin-kindlin-1 axons regenerate through laminin-111 isoform and tenascin-C positive tissue. (**A**) The bridge that develops over lesions is mostly derived from connective and/or meningeal tissue and is therefore GFAP-ve and tenascin+ve laminin+ve. Tangled axons and axons crossing the lesion from α9-kindlin group are seen inside the connective tissue bridge. Scale bar: 200 μm. (**B**) The detailed image shows the bridge region with growing axons with morphology typical for regenerates, and in some cases also with the retracting bulb. Scale bar: 50 μm. (**C**) Axon growth in the α9-kindlin group through connective tissue strands that are tenascin-C stained. Many axons with a typical regenerated morphology are seen within the strands, then entering CNS tissue rostral to the lesion on the left. Scale bar: 200 μm.

In summary, within the lesion axons expressing α9-V5 and kindlin-1 appeared to regenerate preferentially through connective tissue structures that were positive for laminin and tenascin-C, and were then able to re-enter and grow within the tenascin-C-expressing CNS tissue. Axons expressing kindlin-1 alone contain activated forms of integrins expressed endogenously by sensory neurons (α4,5,6,7). These integrins bind to laminin and fibronectin receptors. This means that these axons grew where laminin is present, but did not re-enter the laminin -ve CNS tissue.

### Completeness of dorsal column crush

To verify the completeness of the lesion in our dorsal column crush model, we used 7T MRI. All dissected spinal cords were imaged as sagittal and transversal sections (Supplementary Fig S2). The sections were then compared with the rat anatomical atlas to exclude lesions that were too wide, too deep, or too small. However, no animal was excluded from the study based on the MRI results. Staining of the sensory nuclei further allowed us to demonstrate lesion completeness, as regenerating axons are excluded (Fig. 4E) whereas non-lesioned axons innervate the nuclei ^12,16^. Thus, if V5- or GFP-positive axons were observed in the sensory nuclei, the lesion would be considered incomplete.

**Supplementary Figure S2.**
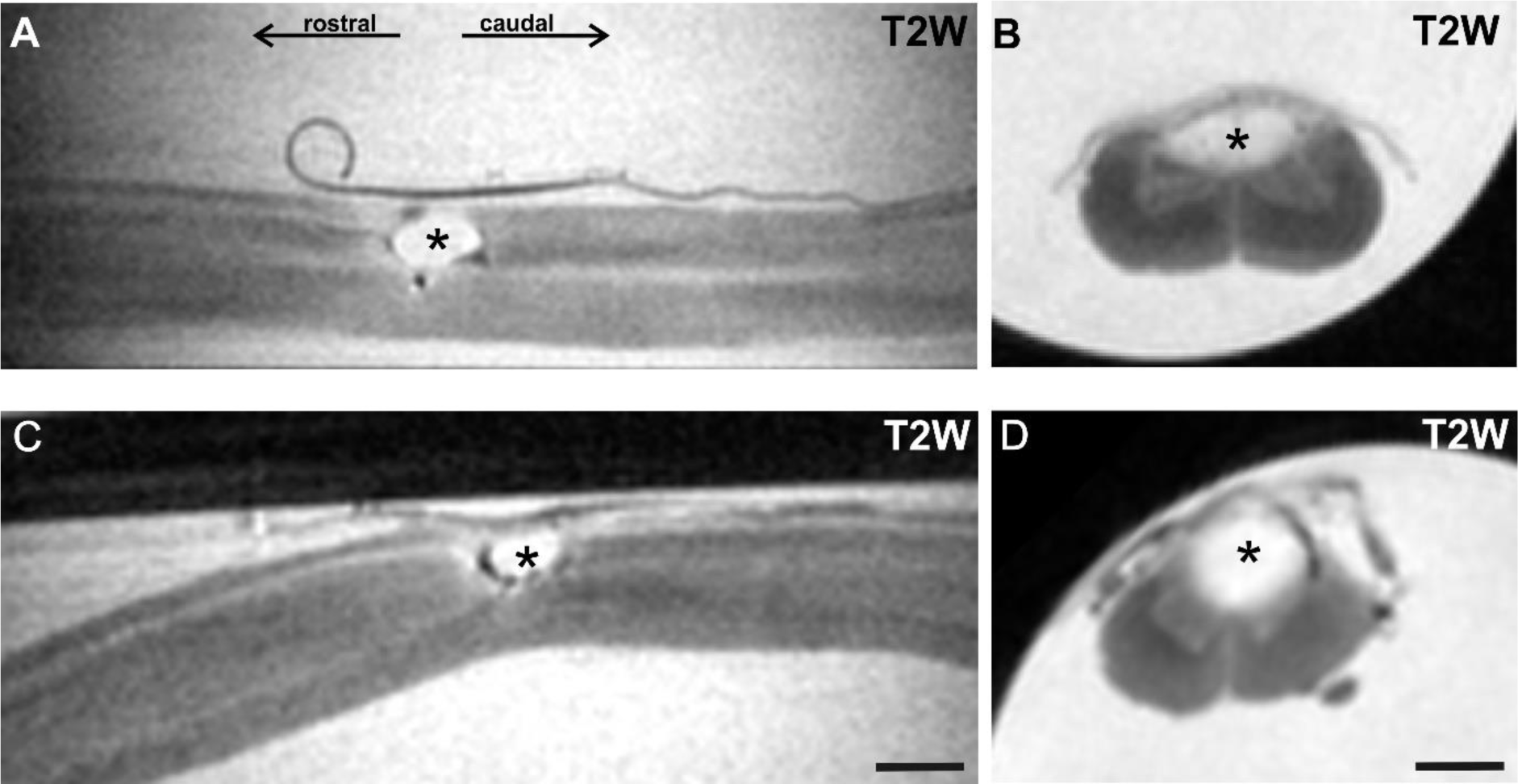
Completeness of dorsal column crush. MRI T2-weighted images in longitudinal (**A, C**) and transverse (**B, D**) orientation to show a typical lesion. These images were taken from spinal cords collected from animals after fixation at week 13. The stars indicate the lesion. (**A, B**) show the C4 lesion. (**C, D**) show the Th10 lesion. Scale bar: 600 μm

### Regenerating axons make a functional synapse above the lesion

To assess functional connectivity over the lesion, spinal cFOS expression was visualised after electrical stimulation of the median and sciatic nerves. cFOS belongs to the immediate early genes and is a well-established marker of transcription induced by neuronal activity (Fig. 7A-C).

**Figure 7.**
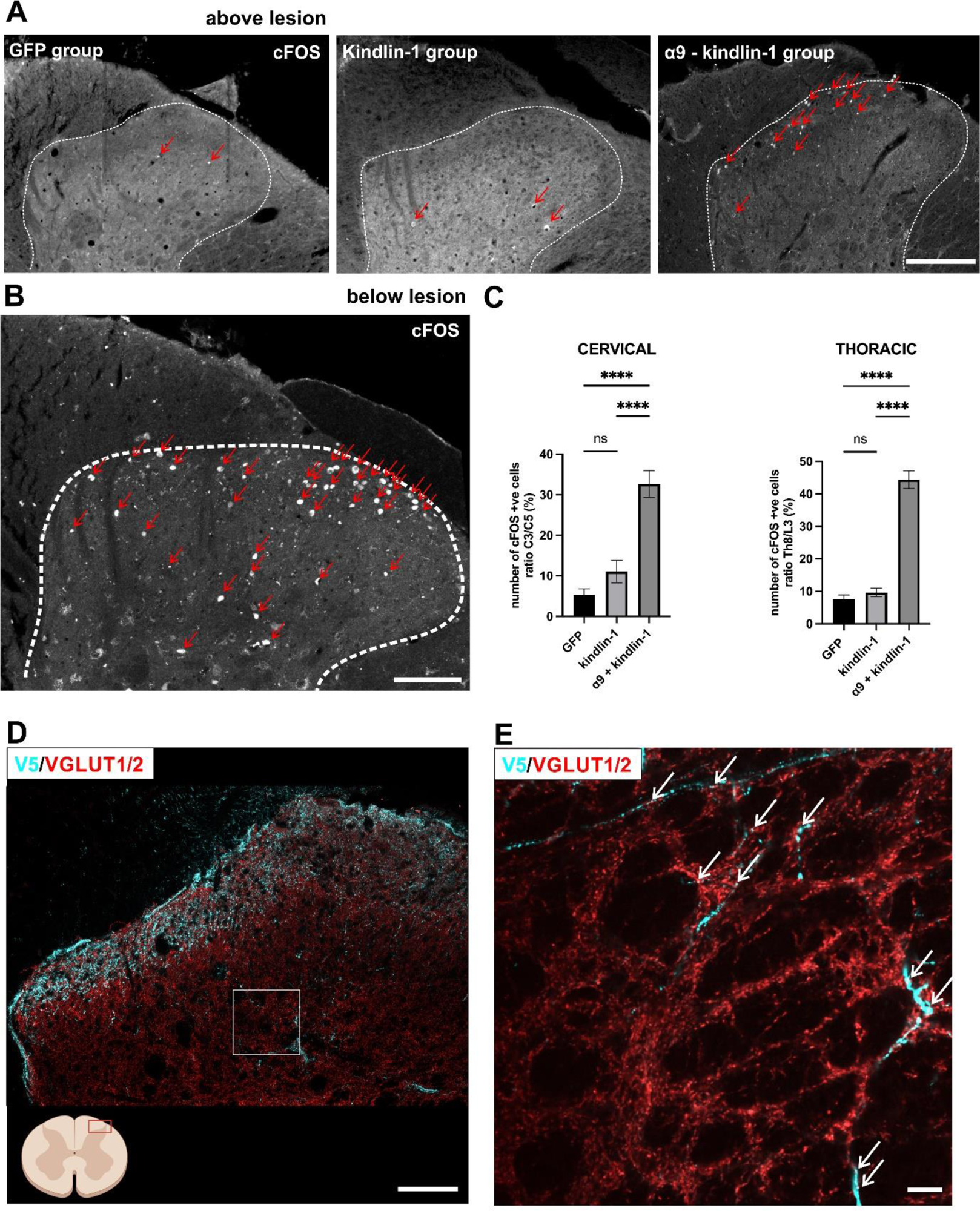
Regenerating axons form a functional synapse above the lesion. (**A, B**) Staining of cFos from the dorsal horn of the spinal cord (white dotted lines delineating the dorsal horn border) after peripheral nerve stimulation. (**B**) Stimulation activates many neurons (red arrows) bellow the lesion. Scale bar: 100 μm. (**A**) 2 segments rostral to the lesion few neurons are activated in the GFP and kindlin-1 groups (red arrows), but many more are activated in the α9-kindlin group (red arrows) indicating that functional connections have been made. Scale bar: 200 μm. (**C**) Quantification of (**A, B**). The number of cFos positive cells is expressed as the ratio of positive cells bellow and above the cervical and thoracic lesion. Data show mean ± SEM (n= 3 animals per group). ns p ≥ 0.05, **** p < 0.0001, one-way ANOVA, Tukey’s multiple comparisons test. (**D**) Representative confocal images showing the dorsal horn 1cm rostral to the lesion from the α9-kindlin group. Blue V5-positive axons are seen extending into the grey matter showing swellings that also red stain for VGLUT1/2. Scale bar: 100 μm. (**E**) Super resolution single-plane images of the dorsal horn detail (indicated by the white rectangle in (**D**) showing blue V5-positive axons growing into the grey matter with swellings, which also stain red for VGLUT1/2. Scale bar: 20 μm.

To quantify the total number of cFOS+ nuclei in the dorsal horns below (C6 or L1 level) and above the lesion (C2 and Th8 level), the Cell Counter plugin in ImageJ ^17^ was used. For each animal, three adjacent sections below and above the lesion were analysed, and then the mean of each subset was calculated. Results are presented as a percentage ratio of the number of cFOS-positive cells above the lesion to the number of cFOS-positive cells below the lesion. A higher percentage of cFOS-positive cells was observed in the integrin α9 – kindlin-1 group (32.656 ± 3.305 % after cervical SCI and 44.380 ± 2.684 % after thoracic SCI) compared to the GFP (5.369 ± 1.451 % after cervical SCI and 7.754 ± 1.334 % after thoracic SCI) and kindlin-1 (11.084 ± 2.759 % after cervical SCI and 9.650 ± 1.313 % after thoracic SCI) groups (Fig. 7A,C). The differences between α9 – kindlin-1 and GFP groups were significant (P<0.0001). No significance was observed between the GFP and kindlin-1 groups in either cohort.

Using super resolution, V5-positive sensory axon terminals were observed colocalizing with VGLUT1/2 puncta in spinal coronal section taken above the lesion, indicating functional synaptic connectivity (Fig. 7D, E). These data show that regenerated axons in the α9 – kindlin-1 group were able to establish functional synapses rostral to the lesions.

### α9-K1 restored sensory functions

To examine recovery of sensory behaviour, forelimb and hindlimb function tests were used in animals with C4 lesions/cervical DRG injections and T10 lesions/DRG lumbar injections respectively. A soft mechanical pressure (Von Frey) test, a heat test (Plantar/Hargreaves) and a tape removal test were used (Fig. 8). The tests started 2 weeks before surgery (pre-test period) to obtain reference values for healthy animals and also to allow the animals to learn the required task. The week after surgery represented a break in testing to allow the animals to adapt to the new pathophysiological conditions. From the second week of viral expression, animals were tested every other week for 12 weeks.

**Figure 8.**
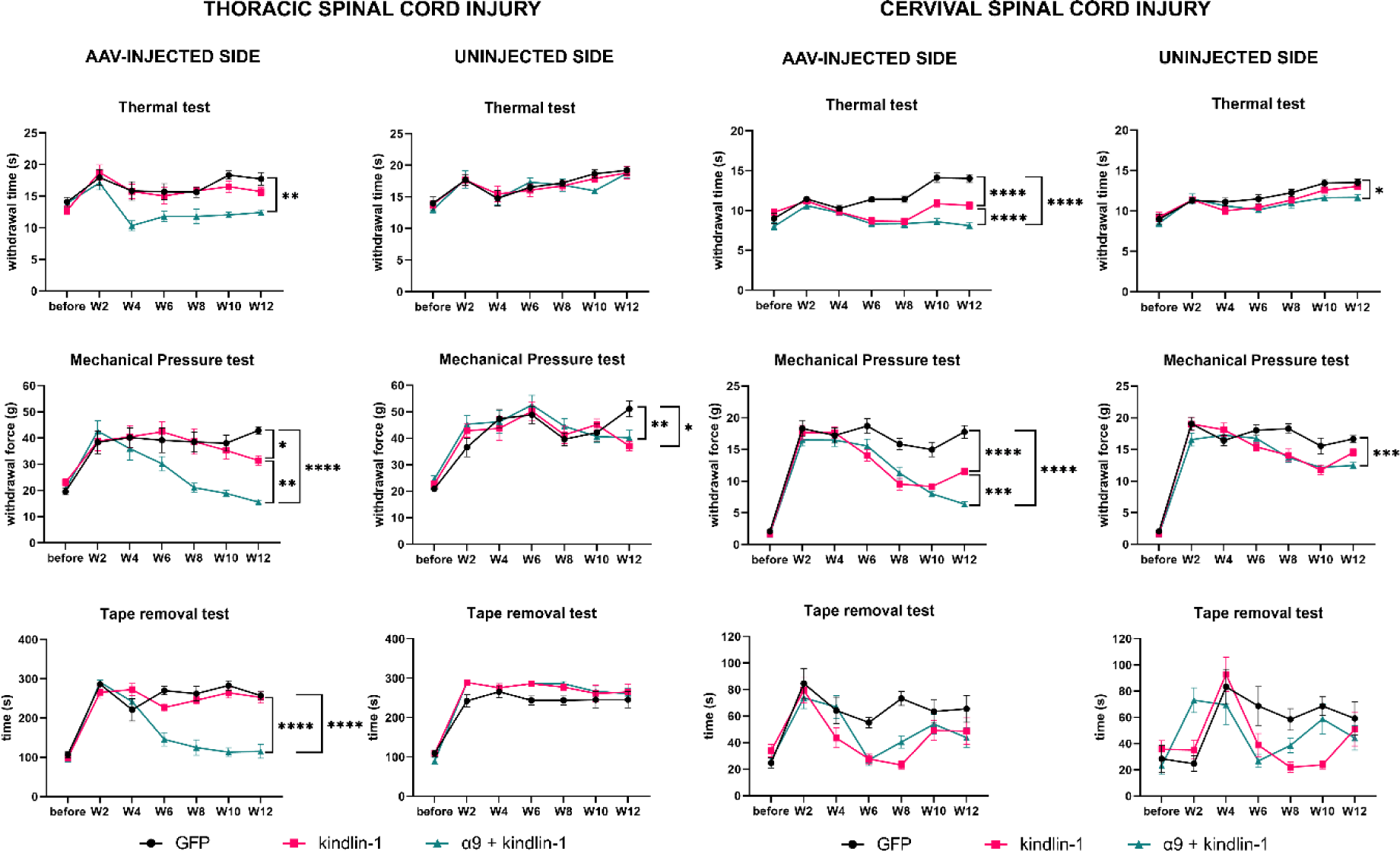
α9-K1 restored sensory functions. Results of sensory tests in animals that received a Th10 lesion and injection into L4,5 DRGs (**A**) a and in animals with C4 lesion and injection into C6,7 DRGs (**B**) After the thoracic lesions, there was a recovery of heat sensation, pressure sensation and removal of the tape only in the α9-kindlin group and only on the treated side. For the cervical lesions some recovery also occurred in the kindlin-1 group. Data showed mean ± SEM (n= 10-12 animals per groups). ns p ≥ 0.05, *p < 0.05, **p < 0.01, ***p < 0.001, **** p < 0.0001, two-way ANOVA, Tukey’s multiple comparisons test.

During the first testing week, the experimental left paw and the internal control right paw showed a significant sensory deficit compared to the reference values after both levels of injury, cervical and thoracic. The sensory deficit was manifested by a greater force required to elicit withdrawal during the mechanical pressure test, a longer withdrawal time during the heat test and a longer delay time with tape during the tape removal test.

In the mechanical pressure test, the integrin α9-kindlin-1 group began to show significant recovery compared to the GFP group from week 6 in the cervical SCI cohort (p = 0.0287, 2way-ANOVA, n =11-12) and from week 8 in the thoracic SCI cohort (p = 0.0006, 2way-ANOVA, n =10-12). In the cervical SCI cohort, the kindlin-1 only group also showed significant but lesser recovery compared to the GFP group (p = 0.0007, 2way-ANOVA, n =11-12) starting at week 6. The other 2 groups (GFP and kindlin-1 only) in the thoracic SCI cohort showed no significant recovery.

In the heat test, both groups, kindlin-1 only and integrin α9 – kindlin-1 showed significant recovery (p < 0.0001, 2way-ANOVA, n = 11-12 per group) compared to the GFP group in the cervical SCI cohort. Significant differences began at week 6. After thoracic SCI, only the integrin α9 – kindlin-1 group showed significant recovery compared to the GFP control group (p = 0.0002, 2way-ANOVA, n = 10-12) from week 4 onwards but there was no recovery in the other groups.

In the tape removal test, both groups, kindlin-1 only (p = 0.0162, 2way-Anova, n = 10-12) and integrin α9 – kindlin-1 (p = 0.0107, 2way-Anova, n = 10-12) showed significant recovery compared to the GFP group in the cervical SCI cohort. Significant recovery was observed from week 6, however, by week 10 the significant difference disappeared, which we attribute to the animals learning the task and rapidly removing the tape guided by vision instead of sensory perception. After thoracic SCI, significant recovery was observed only in the integrin α9 – kindlin-1 group (p < 0.0001, 2way-Anova, n = 10-12) compared to the GFP control group. In contrast to the conditions after cervical SCI, the kindlin-1 only group showed no significant recovery after thoracic SCI.

In summary, these results indicate that the integrin α9-kindlin-1 group exhibited superior sensory recovery in the mechanical pressure and heat tests in both cohorts. They also performed well in the tape removal test after thoracic SCI. Our results suggest that tape removal test after cervical injury is not ideal for long-term studies such as this, as the animals likely saw the tape on their forepaw before they felt it.

## Discussion

This study has demonstrated that α9-K1 transduced neurons regenerated their neurons vigorously through the largely fibroblastic environment of the lesion core of the damaged rat spinal cord. The axons then progressed from this environment back into spinal cord CNS tissue, where they regenerated up to the level of the medulla, a distance of greater than 4 cm from thoracic lesions. Many axonal branches grew into the dorsal horn grey matter, where synapses were evident. On stimulation neurons in the dorsal part of the cord upregulated cFos, indicating that functional connections were formed. Full behavioural recovery was seen for light touch, heat and tape removal.

### Mechanisms of axon regeneration

DRG sensory neurons are regeneration-capable, as evidenced by their ability to regenerate axons within the PNS. Extensive regeneration of the axons in the current study suggests that α9-K1 transduction activates mechanisms that allow PNS regeneration, and enable regeneration in the CNS environment. Sensory regeneration in the PNS is associated with upregulation of the RAGs programme of gene expression ^1^. Several of the molecules expressed in the RAGs programme facilitate successful regeneration, so regeneration without expression of the RAGs programmed is unlikely. Transduction of sensory neurons with α9-K1, even without axotomy, leads to expression of a RAGs programme, priming the neurons for successful regeneration^13^. However, expression of the RAGs programme by itself is not sufficient to enable long-distance spinal sensory regeneration. Crush of peripheral nerves prior to lesion of the central sensory branch upregulates the RAGs programme and leads to increased local sprouting of the cut axons, but not long-distance growth ^18^. An additional element is therefore required. Cell migration events depend on cell surface growth-promoting receptors that match to ligands in the environment ^8^. This receptor-ligand interaction leads to signalling, increased cytoskeletal dynamics linked to focal adhesions, leading in turn to mechanical traction and migration ^19^. The main migration-inducing cell surface adhesion molecules are integrins. Adult DRG neurons express several integrins that bind to fibronectin and laminin (α3β1, α4β1, α5β1 α6β1 and α7β1), with the laminin receptor α7β1 being the main contributor to axon regeneration in peripheral nerves ^20,21^. However, laminin and fibronectin in the acutely injured cord are found on fibroblastic cells in the lesion core, around blood vessels and on the meninges, not in the CNS tissue ^22,23^. The integrin ligands in reactive CNS tissue are tenascin-C and osteopontin, which are not partners for the integrins expressed in adult DRGs. The main migration-inducing integrin for tenascin-C and osteopontin is α9β1, and it is therefore not surprising that α9-K1 transduction enables axon regeneration ^24^. Expression of kindlin-1 alone will activate the integrins endogenously expressed by DRG neurons, which bind to laminin and fibronectin. Therefore kindlin-1 alone allowed sensory axons to regenerate into the lesion core region, but not into CNS tissue which contains tenascin-C, for which the sensory axons lack an integrin receptor, but very little laminin. Regeneration into the lesion and on into CNS tissue was only seen when neurons were transduced with α9-K1 to provide an activated form of this tenascin-C osteopontin receptor. Unlesioned axons below the lesion in α9-K1 transduced animals mostly expressed both α9 and K1, but some were positive for only one of these. Above the lesion we only saw axons that expressed both α9 and K1, showing that this combination is needed for successful CNS regeneration. However, in animals with thoracic lesions, we observed a diminishing V5 signal in the medulla, which may indicate that integrin transport becomes weaker with distance.

### Pathway of axon regeneration

Axons regenerating through the spinal lesion were found associated with strands and bridges of GFAP negative tissue, which has been shown to contain cells of meningeal, vascular, perivascular and fibroblastic origin. These cells expressed laminin and tenascin-C, and therefore provided a matched growth substrate for endogenously-expressed laminin-binding integrins activated by kindlin-1 and for neurons expressing α9. At the rostral interface between lesion core and CNS tissue, there was usually a region in which axons grew chaotically indicating exploratory behaviour in a disrupted tissue, after which their growth was straight and direct. Within the cord rostral to the lesion regenerating axons in the α9-K1 group were found particularly at the boundary between the dorsal columns and the dorsal horn, with a few axons growing in a wandering fashion through white matter. This differs from the normal pathway of sensory axons, and also differs from the path through white matter that was taken by axons regenerating after α9-K1 and a dorsal root crush ^12^. We assume that the pathway disruption associated with growth through the lesion cause axons to search for a permissive path, although why the white matter/grey matter boundary should provide this is not obvious. As in previous studies, the regenerating sensory axons did not penetrate the medullary sensory nuclei ^12,25^. This is due to the dense CSPG-rich nets found in these nuclei. We saw differences in regeneration within the cord in animals with cervical lesions and cervical DRG α9-K1 treatment. More axons in this group continue to grow beside the spinal cord in meninges rather that re-entering CNS tissue. In the cervical SCI, the axon entry zone was much closer to the lesion itself, which could cause the axons to grow more in the meninges and not re-enter the grey matter as in the thoracic SCI. Therefore, preventing waste of regenerating axons that choose to grow in the meninges rather than spinal cord will be an issue for further development.

### Regeneration vs. sparing

In this study, and in our previous study using dorsal root crush rather than spinal lesions ^12^, around 1000 axons regenerated 2-5cm from lesions to the hindbrain. Several data show that this is regeneration rather than unlesioned axons. 1) The axons in the lesion pass through GFAP-ve fibroblastic tissue; unlesioned axons would be surrounded by CNS glia 2) Axons rostral to the lesion follow a different pathway to uncut axons, 3) The regenerated axons avoid the sensory nuclei; there are no axons innervating the sensory nuclei (if present these would be unlesioned axons), 4) The controls are very clear, no axons past the lesion in the GFP group, axons only in the laminin+ve fibroblastic tissue in the kindin-1 group.

### Sensory recovery and synapse formation

Many processes were seen growing into the dorsal horns, taking a course perpendicular to the ascending axons. VGLUT1/2 +ve swellings were seen on these processes, indicating synapse formation. We tested for the ability of these synapses to stimulate neurons in the cord by observing upregulation of cFos in propriospinal neurons after stimulation of the median/sciatic nerve, the peripheral branch of the DRG injected with α9-K1. In α9-K1-treated animals the number of cFos neurons after stimulation was much greater than in controls, indicating connections between regenerated axons and cord neurons. This was reflected in sensory recovery tests. Particularly in the lumbar injected α9-K1 group, we saw eventual complete recovery in fine touch, heat and tape removal tasks. The time course of recovery was similar to the progression of axon regeneration in this and our previous experiment ^12,13^. Recovery in the tape removal task is particularly informative. Animals do not appear to see tape on their hind paws, unlike the forepaws where tape removal is often triggered by visual recognition. Instead hind paw tape removal appears to be triggered by sensory recognition. In order to sense that there is tape on the hind paw, sensation must reach the brain which then executes the removal process. This task therefore shows that regenerating axons carry information to the brain. It is interesting that this occurs in the absence of re-innervation of the medullary sensory nuclei. Presumably sensory information is relayed to the brain from propriospinal neurons that were stimulated by regenerated sensory inputs to the dorsal horn.

The strategy of using an activated integrin to induce sensory regeneration has enabled almost complete reconstruction of the spinal sensory pathway. The lesion length in human spinal cord injuries varies between 1 and 7cm ^26^, so it is important that in the present study axons were able to regenerate for a length that could enable growth across a human injury. The regeneration index, comparing axon numbers below and above the lesion indicates that 50% of axons regenerated through the lesion area, and once through the lesion the number of axons did not decline much up to the high cervical cord. However complete reconstruction was not achieved because innervation of the medullary sensory nuclei did not occur. Methods of enabling re-innervation of these nuclei have been identified using chondroitinase digestion and expression of NT3 ^25,27^, and these could be applied to bring about a complete tract reconstruction. However, despite lack of innervation of the sensory nuclei functional recovery was almost complete, including tape removal which requires that sensory information must reach the brain. If α9-K1 could be delivered to descending spinal axons they would probably enable regeneration. However, this repair strategy cannot at present be transferred directly to descending motor pathways because in these highly polarized neurons integrins are restricted to the somatodendritic domain and excluded from axons ^28,29^. Strategies to enable integrin transport to motor axons have been identified, which may make it possible to reconstruct motor pathways ^30,31^.

## Data availability

Data available on the request from the corresponding author.

## Acknowledgement

Microscopy was done at the Microscopy Service Centre of the Institute of Experimental Medicine CAS supported by the Ministry of Education, Youth and Sports of the Czech Republic (LM2023050 Czech-Bioimaging). MRI was done at the core facility Center for Advanced Preclinical Imaging (CAPI) Charles University in Prague supported by MEYS CR (LM2023050 Czech-Bioimaging) and by European Regional Development Fund (Project No. CZ.02.1.01/0.0/0.0/18_046/0016045).

